# *SAMHD1* Knockout iPSC model enables high lenti-viral transduction in myeloid cell types

**DOI:** 10.1101/2025.02.04.636295

**Authors:** Huinan Li, Maliha Afroze, Gunisha Arora, Scot Federman, Kaivalya Shevade, Yeqing Angela Yang, Phuong Nguyen, Rustam Esanov, Laralynne Przybyla, Adam Litterman, Shawn Shafer

## Abstract

Recent advances in functional genomics tools have ushered in a new era of genetic editing to identify molecular pathways relevant to developmental and disease biology. However, limited model systems are available that adequately mimic cell states and phenotypes associated with human disease pathways. Here, we quantitatively analyzed the founder population bottleneck effect and demonstrated how the population changes from induced pluripotent stem cells (iPSCs) to hematopoietic stem cells and to the final induced macrophage population. We then engineered *SAMHD1* knockout (KO) iPSC and characterized the iPSC line with RNA Seq, and induced macrophages from two distinct protocols with functional analysis. We then generated *SAMHD1* KO CRISPR-dCAS9 KRAB iPSC through lenti-viral transduction aiming to increase the efficiency of lentiviral mediated gene transfer. We demonstrated increased lenti-viral transduction efficiency in induced macrophage, as well as microglia induced with two distinct protocols. This model allows for efficient gene knock down, as well as large-scale functional genomics screens in mature iPSC-derived macrophages or microglia with applications in innate immunity and chronic inflammatory disease biology. These experiments highlight the broad applicability of this platform for disease-relevant target identification and may improve our ability to run large-scale screens in iPSC-derived myeloid model systems.

## Introduction

Macrophages and their tissue-resident counterparts such as microglia in the brain, perform essential roles during development, homeostasis, and immunological disorder, and are key cellular players in disease such as neurological disorders in the central nervous system^1,2^. The states of macrophage and microglia are altered in response to specific signaling pathways during development, disease, and in situations of acute or chronic inflammation^3,4^. In order to model human macrophages *in vitro*, primary cell cultures and humanized rodent models have been widely utilized to gain new insights into the initiation and progression of disease states^2,5^. However, such models often pose challenges when it comes to genetic editing process as well as scalability. iPSC derived macrophages and microglia have addressed several of these challenges, including high efficiency of induction, faithful recapitulation of the innate gene signature, and response to physiologically relevant insults, which provides new avenues to model human disorders. Recently, with advances in CRISPR technology, the capability and scalability of editing approaches have been greatly extended allowing for modification of the human genome, epigenome and transcriptome, followed by cell type-specific phenotypic readouts leading to a more comprehensive understanding of specific gene function^6^. This allows for functional phenotypic analyses to be conducted, but other challenges remain. Macrophages and microglia are known to be resistant toward many forms of DNA delivery including electroporation, transfection, and viral transduction^7^. For this reason, iPSC-induced macrophage and microglia are difficult to engineer to large scale particularly for introducing libraries of hundreds or even thousands of perturbations at a time.

Most iPSC cytokine-induced differentiation protocols don’t investigate how the population changes from the founder iPSC population to the progenitor or mature differentiated states, but in many cases the differentiated population is selected or sorted out resulting in a population bottlenecking over the course of the differentiation trajectory^8–10^. Using a transcription factor overexpression approach to deterministically drive differentiation provides an alternative route for generating a uniform population for functional genomic analysis. However, iPSC line engineered to overexpress transcription factors are difficult to generate and time consuming^11,12^. In order to use a conventional cytokine-induced macrophage differentiation protocol in the context of functional genomic perturbation, we sought to develop a lentiviral strategy that improves macrophage and microglia viral infection efficiency. Human sterile alpha motif and HD domain containing protein 1 (*SAMHD1*) degrades deoxynucleotide triphosphate (dNTP) into the 2′-deoxynucleoside (dN) and triphosphate subunits, maintaining the balance of intercellular dNTP pools and maintaining nucleotide homeostasis. In different cell types, SAMHD1 acts to either promote (e.g. dividing T cells^13,14^) or restrict (e.g. nondividing macrophages^15^) viral replication by maintaining homeostasis of the dNTPs. Previously, it was discovered that SAMHD1 activity protects monocytes and macrophages from viral infection by degrading dNTPs^16,17^. Inhibition of *SAMHD1* expression by RNAi or over-expression of a known repressor of SAMHD1 function viral protein X (VPX) has been shown to increase efficiency of viral transduction in monocytes and macrophages^18^. THP-1 human monocytes (or THP1 differentiated macrophages) are widely used to study DNA sensing activity. This cell type expresses double-strand DNA sensors including cGAS, IFI-16, DDX41 and LRRFIP1 *etc*. A widely used THP-1 *SAMHD1* KO monocyte cell line^19^ provides a useful model to investigate IRF and NF-κB pathways, discover interaction between *SAMHD1* and other potential signaling proteins and identify the role of SAMHD1 in innate immunity, tumorigenesis or viral replication^20^. However, to our knowledge, there is no published approach for generating *SAMHD1* KO iPSC induced macrophages.

We conducted a quantitative analysis of cell population diversity during induction to macrophages using two distinct protocols. This was done to examine the extent of population bottlenecks throughout iPSC-derived macrophages (iMacrophages) differentiation. Through barcode labeling founder iPSC population, we tracked the founder cell diversity all the way to iMacrophages. We identified significant bottleneck effects in the iPSC diversity during iMacrophages induction process. This bottleneck effect encouraged us to explore alternative options that would allow for functional genomic studies in iMacrophages through lentiviral transduction. We then engineered an iPSC line with *SAMHD1* gene knockout. The loss of *SAMHD1* has little effect on iPSC health, and these cells can be induced into macrophages and microglia with multiple induction protocols. RNAseq and functional tests on the differentiated populations indicate that knocking out *SAMHD1* also has little effect on the overall health of the induced cells including iMacrophages and iMicroglia. In addition, we engineered *SAMHD1* KO dCAS9 KRAB CRISPRi iPSC line. We demonstrate that iMacrophages and iMicroglia have dramatic increase in transduction efficiency with various lentiviral cargo types. We believe this model can be used for CRISPR functional genomic analysis, including introducing point mutation, knock out and knock in of specific genes for functional genetic studies, even for large scale CRISPR screens.

## Results

### Quantification of population diversity during macrophage differentiation

To understand the ability of differentiated macrophages to be used for functional genomics screening, we employed two different differentiation protocols. The first was a published protocol for obtaining iMacrophages^10^ (hereafter iMacrophages) and the second used a commercially available kit ^9,21^ (StemCell Technologies, hereafter iMacrophages ST). We compared two distinct differentiation protocols for macrophage differentiation from iPSCs (Fig. 1a) and analyzed their effects on gene expression and population bottlenecking upon transduction with a pooled viral library. To confirm that the two induction protocols generated the desired cell types, we performed bulk RNA sequencing and differential expression analysis (Fig. 1b). Based on gene signatures, we were able to distinguish iPSC, iPSC-derived hematopoietic progenitor cells (iHPCs) and iMacrophages. iMacrophages from these two protocols were very similar as confirmed with principal component analysis (Fig. 1c). In the second protocol ^9,21^, iMacrophage precursors (iMacrophage Pre) were transcriptionally like fully differentiated iMacrophages (Figure 1b), consistent with the original description of this protocol^10^. To quantitatively understand the founder cell population identity changes throughout the whole induction, we performed these iMacrophages protocols using barcoded iPSC populations. iPSCs were infected with virus containing unique barcodes, then these cells were selected through puromycin and continued to proliferate until there were about 80% RFP (red fluorescent protein) positive. These populations of cells were divided into 3 groups, each with equal amount of iPSCs as founder cell population to ensure adequate representation of barcodes throughout the inductions (Fig. 1d). One third of iPSCs were saved for sequencing as the initial founder cell population identity (iPSC); the second third of iPSCs were induced with a previously described protocol^10^. The final third of iPSCs were induced into macrophage with another protocol previously described^9,21^. According to the final data analysis, we were able to recover 260,050 unique barcodes in the founder population iPSCs, after induced into iMacrophages with these two protocols, the barcode diversity decreased to 40,810 for iMac population and to 14,998 for iMac ST population. We also confirmed that there was overrepresentation of specific barcodes with a Shannon evenness index for iPSC population 0.909, iMacrophages 0.763 and iMacrophages ST 0.570, indicating founder population drift phenomenon. This indicates that there are strong bottlenecks limiting the iPSC diversity upon differentiation with either protocol.

**Figure 1.**
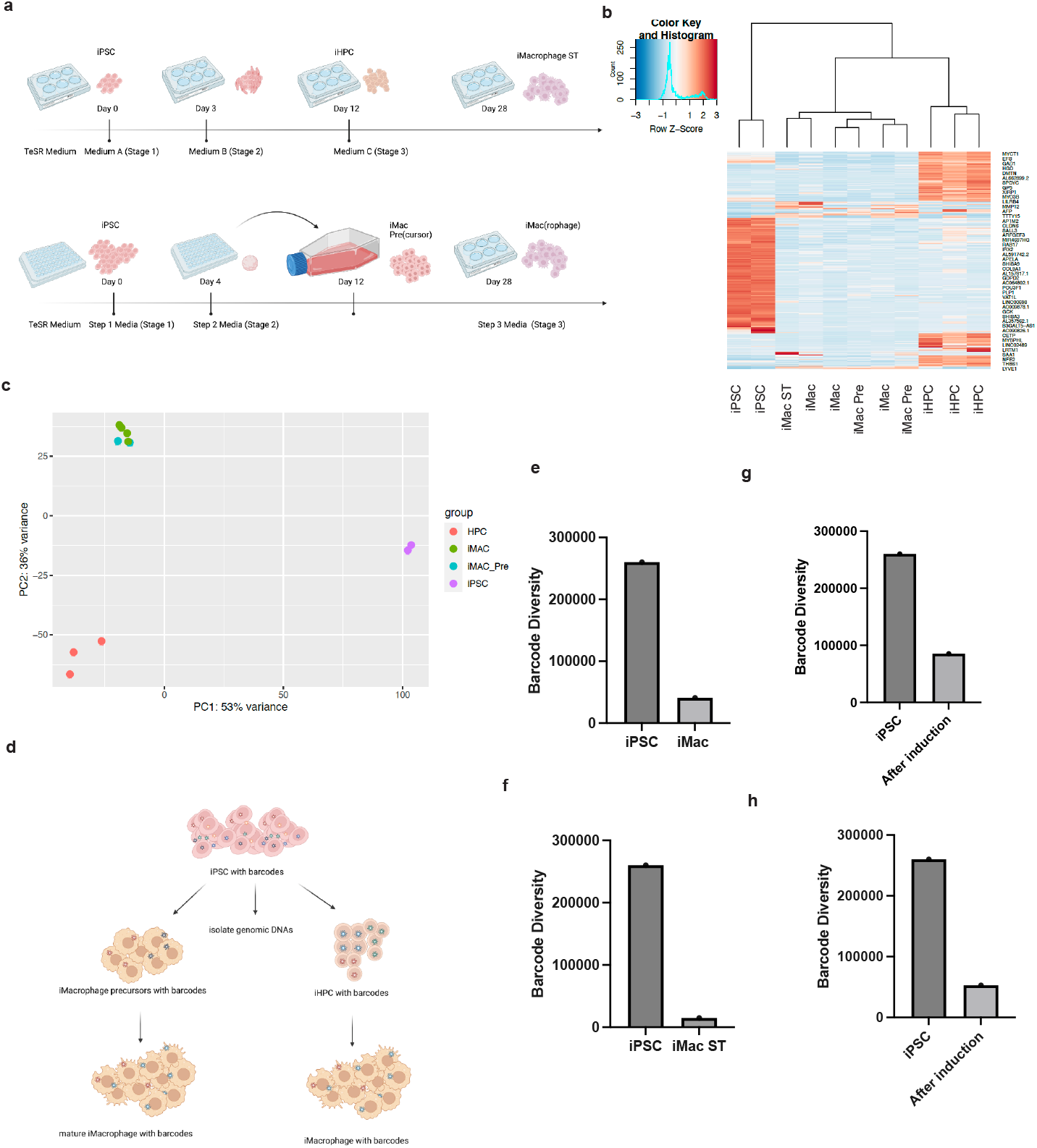
Founder Cell population Analysis of iMacrophages across two distinct protocols. a. Schematic illustration of both iMacrophage induction protocols. Upper, iPSCs cultured in mTeSR1 media were induced into iHPC first with addition of media A and media B before induced into iMacrophage (iMac ST) with media IMDM with FBS, supplemented with MCSF, GMCSF and IL-3; lower, E8 Flex maintained iPSCs were cultured in E8 Flex media supplemented with BMP4, VEGF,SCF in 384 well round bottom plate for 4 days before induced into immature macrophage with StemPro34, supplemented with IL-3, MCSF,Glutamax, β-mercaptoethanol; the iMac Pre(cursors) were then transferred to 6 well plate into iMacrophage (iMac) with RPMI, FBS, Glutamax, MCSF, b. RNA sequencing data cluster analysis indicates duplicated samples in each stage express specific marker genes, c. Principle Component Analysis indicate iPSCs clusters together, away from iHPC, and iMac of both induction cluster together, as well as iMac precursors, d. Schematic illustration of sampling iPSC with barcodes across both inductions, e. iPSC, iMac and iMac ST three populations of the individual barcode counts.

### Generation and characterization of SAMHD1 KO iPSCs and differentiated cells

*SAMHD1* is a non-essential gene and has been demonstrated to inhibit viral infection in macrophages, so we hypothesized that *SAMHD1* mutant macrophages would be more permissive of lentiviral gene transfer^22^. To generate a *SAMHD1* knock out iPSC cell line, we performed a Cas9 RNP electroporation with 3 synthetic single guide RNAs targeting different regions of *SAMHD1* exon1 into S2-1 iPS cells (Figure 2a). Gel electrophoresis analysis of a PCR amplicon of this exon confirmed the generation of a deletion that reduced the amplicon size by ∼100 base pairs (Figure 2b). Next, we used bulk RNA sequencing to examine the effect of *SAMHD1* knockout on gene expression in iPSC and differentiated iMacrophages. (Figure 2c). In both iPSC and iMacrophages, the gene expression profiles were largely similar between WT and *SAMHD1* KO cells (Figure 2 d,e). We also compared *SAMHD1* expression in different cell types from the RNA sequencing data, using *TREM2* expression as a marker of macrophage differentiation. *SAMHD1* was expressed less in iPSC and iHPC stages, with increased expression in macrophage precursors and the highest level of expression in mature macrophages (Figure 2f). Furthermore, we profiled LPS (lipopolysaccharide), a known activator of macrophages^23^. Cytokine profiling of LPS treated *SAMHD1* knockout and WT iMacrophages revealed that among the 48 cytokines we analyzed, only IP-10 had a significantly different response, lower in the *SAMHD1* knockout macrophages (Figure 2g). We then introduced a dCAS9-KRAB-BFP transgene (#188320, Addgene)^24^ by lenti-viral transduction and sorted BFP positive cells to isolate a pool of cells expressing the CRISPRi machinery.

**Figure 2.**
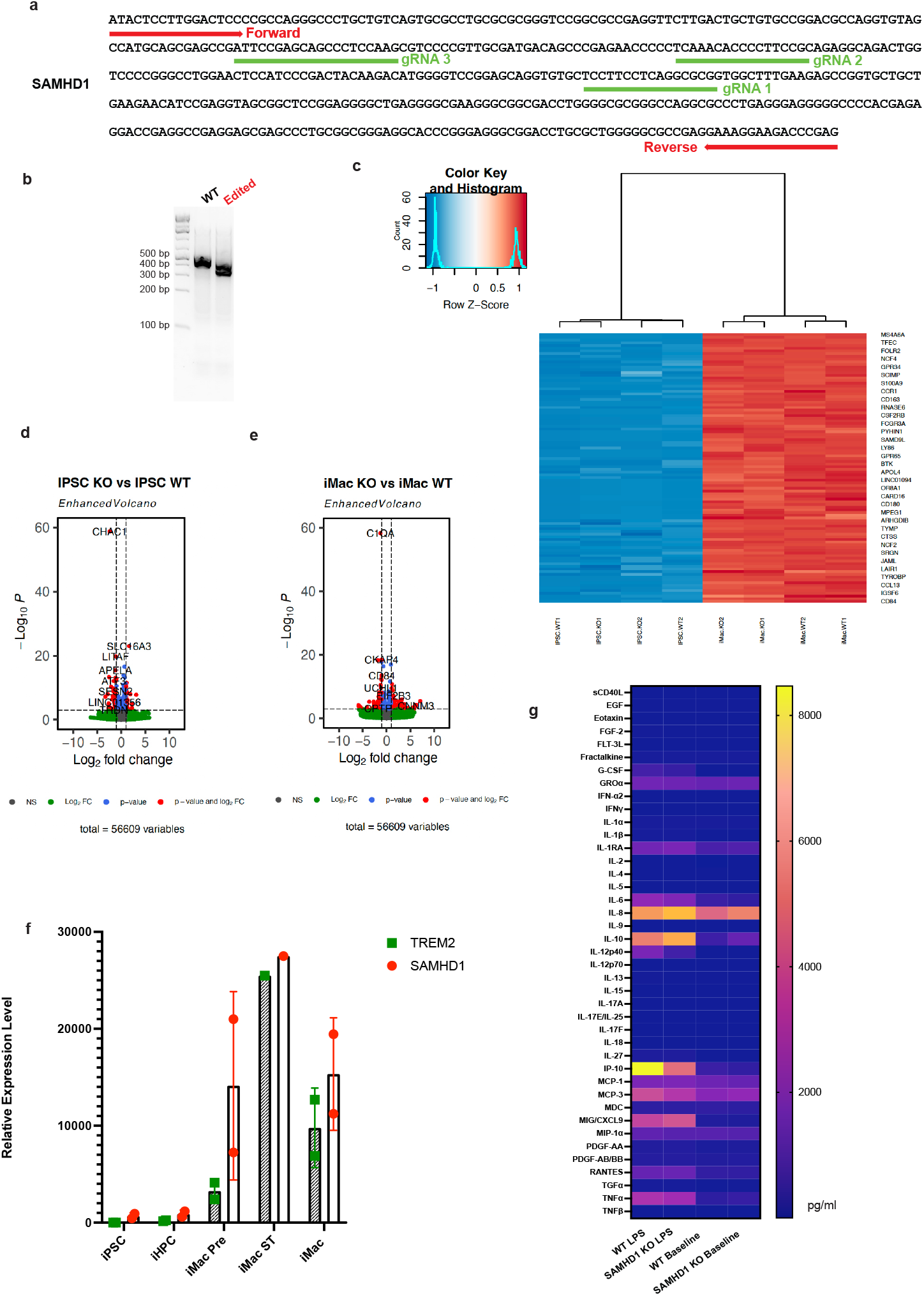
SAMHD1 knockout iPSC line generation and characterization. a. *SAMHD1* gene map sequence with forward and reverse primer sequences, as well as the positions of three guide RNAs, b. PCR gel images indicating the truncation of the gene, c. duplicated iPSCs from two backgrounds and the induced macrophages show little difference, d. volcano plot shows the differentially expressed genes between WT and *SAMHD1* KO iPSCs, e. volcano plot shows the differentially expressed genes between WT and *SAMHD1* KO iMacrophages, f. comparison of *SAMHD1* gene relative expression from RNA sequencing in different cell types, with myeloid lineage marker TREM2, g. cytokine profiling using WT and *SAMHD1* KO iMacrophage prior to LPS treatment and after LPS treatment.

### SAMHD1 KO CRISPRi (dCAS9-KRAB) iPSC induced microglia and macrophage have higher lenti-viral transduction rates

In order to test the effect of *SAMHD1* on different cell types, we induced iPSC with *SAMHD1* KO as well as CRISPRi machinery into myeloid lineage cell types. First, we induced the iPSC into macrophages. iPSCs were first induced into mesodermal lineage and into iHPCs (Hematopoietic Progenitor Cells) in a serum-free and feeder-free condition. These iHPCs express CD34, CD45 and CD43. Then iHPCs were matured into iMacrophages with the addition of cytokines including IL-3, MCSF and GMCSF. To test the ability of these iMacrophages to be transduced upon maturation, we first produced a lentivirus containing a single guide RNA as well as BFP^24^ and transduced *SAMHD1* knockout and WT iMacrophages. One week after transduction we observed 1.37% BFP+ cells in WT iMacrophages versus 3.73% in *SAMHD1* KO iMacrophages (Figure 3b). We observed a more marked difference in transduction efficiency using a commercially available concentrated GFP lentivirus (Millipore) with 0.73% WT iMacrophages exhibiting GFP positivity, while 13.07% of *SAMHD1* KO iMac were GFP+ (Figure 3b).

**Figure 3.**
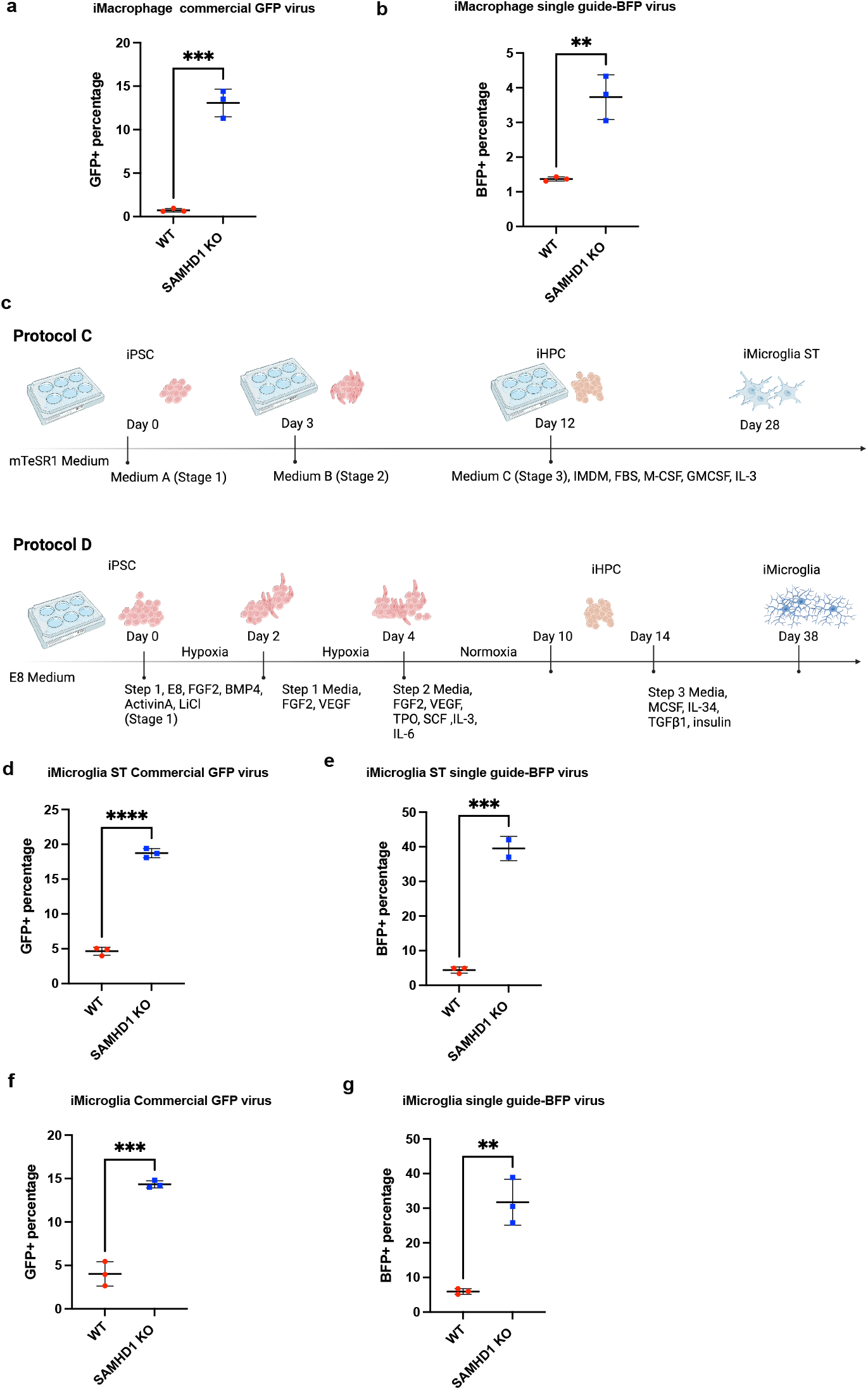
Comparisons between *SAMHD1* knockout CRIPSR dCAS9-KRAB iPSC line induced macrophage and microglia demonstrate increased lenti-viral trandusction efficiency. a. GFP positive cell percentage of iMac of WT and *SAMHD1* KO GFP lentivirus transduction, b. BFP positive cell percentage of iMac of WT and *SAMHD1* KO BFP and CRISPRi single guide lentivirus transduction, c. Schematic drawing of two protocols of iMicroglia induction, d. GFP positive cell percentage of iMicroglia of WT and *SAMHD1* KO GFP lentivirus transduction, e. BFP positive cell percentage of iMiocrglia of WT and *SAMHD1* KO BFP and CRISPRi single guide lentivirus transduction, f. GFP positive cell percentage of iMGL (hypoxia induced microglia) of WT and *SAMHD1* KO GFP lentivirus transduction, g. BFP positive cell percentage of iMGL (hypoxia induced microglia) of WT and *SAMHD1* KO BFP and CRISPRi single guide lentivirus transduction.

We then examined whether the above findings of increased transduction rate in *SAMHD1* KO iMac applied to other phagocytic cell types derived from iPSC. We examined lentiviral transduction rates in three other cell types: iMacrophages ST, iMicroglia derived from a published differentiation protocol ^8^ (iMicroglia) and iMicroglia derived from a commercially available differentiation kit^21^(Stemcell Technologies, hereafter iMicroglia ST). Cells derived from iMicroglia induction became over 80% CD11b and CD45 double positive and over 50% TREM2 positive, reflecting a yolk sac derived microglial cell state^25^. For both iMicroglia and iMicroglia ST we observed strikingly higher transduction rates for *SAMHD1* KO cells with both the guide BFP and GFP lentiviruses (Figure 3 d, e). For example, the increase in transduction in *SAMHD1* KO was nearly 10-fold using the GFP virus in iMicroglia ST with the WT cells having a transduction rate of 4.41% versus 39.5% for *SAMHD1* KO cells Figure 3 f, g). These data indicated that in three different phagocytic cell types derived from a *SAMHD1* KO iPSC cell line using different differentiation protocols, *SAMHD1* KO allows cells to be transduced with markedly higher efficiency.

## Discussion

Macrophages and microglia play essential roles in development, tissue homeostasis, and immunological and neurological disorders^1,3,26^. CRISPR genome editing and screens in rodent macrophages have provided information on genes that are essential for survival and response toward inflammation^27,28^. In addition, human primary cell lines, human immortal cell lines and genetically engineered rodent models have been widely used in the research community and led to informative discoveries^29^. There are also many lines of evidence indicating human and rodent macrophages are vastly different in terms of disease phenotypes^30^. Therefore, *in vitro* models are needed as powerful tools for functional genomics purposes. Human iPSCs can be differentiated into disease-relevant cell types in a large scale^31^. Many cytokine-driven induction protocols recapitulate key features of innate cell populations and allow for human disease relevant biology studies^32^. Alternatively, through overexpressing transcription factors, iPSC can be directly induced into microglia like cells^11,12^. These methods allow for a rapid induction of a homogenous population of microglia cells for large scale functional genomic investigation^11,12^. However, it is known that such transcription factor overexpression strategies require extensive stem cell engineering and characterization, by insertion of more than two gene coding regions. These strategies may therefore be challenging to use with specific iPSC parental lines.

Macrophages and microglia are known to be refractory to commonly employed methods for gene transfer^3^. This characteristic hinders implementation of approaches that require gene manipulation and scalable transduction, including cell type specific response and CRISPR editing as well as large scale CRISPR screens. *SAMHD1* is a key component during viral defense and is a non-essential gene^18,20,22^. It has been demonstrated inhibiting the function of *SAMHD1* can increase viral transduction^14,18,33^.

In this study, we quantitively analyzed the initial founder cell population diversity across the induction with two distinct protocols^9,10,21^, induction from iPSC into macrophages by assigning barcodes to individual iPSC. With the protocol that generated HPCs first, and then matured to macrophages^9,21^, we found that the diversity of the barcodes remained similar until HPC stage, and plummeted to approximately one tenth of the initial iPSC population diversity by the mature macrophage stage. With the protocol that generated immature macrophages first^10^, and then matured to macrophages, we found similar effects, with immature macrophage diversity similar to the original iPSC diversity but dropped to about one fifth of the initial diversity by the mature macrophage stage. This analysis is the first known quantitative analysis tracking barcode diversity during iPSC induction into macrophages and provides quantitative evidence of a loss of library diversity during differentiation. Therefore, functional genomics screens performed by transducing libraries of genetic perturbations into iPSC will have only one tenth of the diversity of the original transduction upon differentiation into macrophages. Our study highlights the need for approaches that bypass this bottleneck effect, which would require much higher screening coverage and could lead to dropouts and false negative results in a screening experiment. Compared with previous literature that used exogenous Vpx addition to degrade *SAMHD1*, our method has less potential for cytotoxicity and phenotypic artifacts caused by the pleiotropic biological effects of Vpx addition^34,35^. We have demonstrated that *SAMHD1* KO iMacrophage generation and iMicroglia generation each using two protocols exhibit increased viral transduction. These engineered cell lines therefore enable transfer of guide RNAs with high efficiency at the macrophage and microglia stage, obviating the need for transduction at the iPSC stage and bypassing this bottleneck effect. This finding was concordant across the four differentiation protocols, indicating that *SAMHD1* KO is a robust method for enabling genetic manipulation of multiple types of iPSC derived phagocytic cells that are refractory to lentiviral transduction. While *SAMHD1* KO had a large effect on the permissivity of phagocytic cells to lentiviral transduction, its overall biological effects were otherwise minimal. By analyzing RNA sequencing and functional cytokine elaboration data from WT and *SAMHD1* KO macrophages, we observed minimal changes to gene expression and functionality, including expression of key markers of macrophage differentiation and elaboration of critical macrophage derived cytokines in response to LPS stimulation. Use of our *SAMHD1* KO approach provides a method for genetic manipulation of a number of human phagocytic cell types, overcoming the difficulty in achieving diverse pools of genetically perturbed cells in these systems. In the future, we anticipate that this approach will empower large scale genetic screens in relevant cellular models of human immunological and neurodegenerative disease.

## Author contributions

H.L., R.E., L.P. and S.S. contributed to the study’s overall conception, design, and interpretation. H.L., L.P. and A.L. co-wrote the manuscript. H.L. created the figures with input from the other authors. H.L. designed and conducted the founder cell population analysis experiment with guidance from L.P. S.F. conducted computational analysis. K.S. and A.Y. conducted RNA sequencing on *SAMHD1* KO and WT iPSC. H.L. induced iMacrophage, iMicroglia from two distinct protocols and compared lentiviral transduction efficiency on *SAMHD1* KO iMacrophage and iMicroglia cells. R.E. designed, generated, and characterized constructs *SAMHD1* KO cell lines. M.A. and R.E. generated *SAMHD1* KO mCherry-dCAS9 iPSC cell line. P.N. made CRISPRi, single guide lentivirus.

## Acknowledgments

We would like to acknowledge Eric Chow for helping to design the barcoding experimental concept.

## Material and Methods

### *SAMHD1* KO and CRISPRi line engineering

The human biological samples were sourced ethically and their research use was in accord with the terms of the informed consent under an IRB/EC approved protocol. To generate SAMHD1 KO iPSC line, gRNA pool from Synthego targeting first exon of SAMHD1 gene (gRNA1: AAAGCCACCGCGCCUGAGGA, gRNA2: UCUGCGGAAGGGGUGUUUGA, gRNA3: CUUGGAGGGCUGCUCGGAAU) was used. Cells were dissociated with Accutase and 500K cells were resuspended in 20ul of P3 solution (Lonza) and 5ul of RNP mix (40pmol of Cas9 and 300pmol of gRNA mix) for nucleofection reaction (Lonza Nucleofector, code CA137). Post nucleofection, cells were seeded onto a geltrex coated plates and incubated in mTeSR1 media in the presence of ROCK inhibitor (10uM) overnight. 3 days after, cells were used for DNA extraction (Lucigen QuickExtract) and PCR to confirm knockout. For PCR, primers spanning gRNA binding cites were designed (Forward: ATACTCCTTGGACTCCCCGC, Reverse: CTCGGGTCTTCCTTTCCTCG) and reaction was performed using Ultra II Q5 DNA polymerase (NEB). PCR products were run on 1% agarose gel for visualization. To introduce CRIPSRi machinery for screening, SAMHD1 KO iPSC line was transduced with a lentivirus encoding for dCas9-KRAB-P2A-mCherry at MOI of 0.1 overnight in the presence of polybrene (1ug/ml). After transduction, iPSCs were expanded and mCherry positive cells were sorted by FACS.

### Barcoding iPSC populations

Barcoding iPSCs were made according to manufacture instruction (CloneTracker XP^TM^ Lentiviral Barcode Libraries). Pilot experiments were conducted to determine cell doubling time and to generate a killing curve. After virus were made with HEK 293 cells, iPSCs were transduced with the library. The second day verified 12% transduction rate. iPSCs were selected with puromycin until over 90% are selected. iPSCs were passaged until reaching around 4.5 millions in total to proceed to induction. These population of cells were divided into 3 groups, each with 1.5 million iPSCs to ensure adequate representation of barcodes throughout the inductions (Fig. 1d). For a total of 1.5 million cells, the first 1.5 million cells were saved for sequencing as the initial founder cell population identity (iPSC); the second 1.5 million cells were induced with a previously described protocol, briefly, iPSC on Day 20, 1.5 million macrophage precursors in T175 flask were harvested again for sampling analysis, and 1.2 million macrophages cultured in a 6 well plate with step 3 media were harvested for sampling analysis; on Day 25, total of 4 million mature macrophages (iMacrophage) were divided into 2 million and 2 million cells for sample analysis. The third 1.5 million cells were induced into macrophage with another protocol previously described. Briefly, on Day 0, 1.5 million iPSCs were cultured with media A in 6 well plates, on Day 2, half media A was changed; on Day 3, media B were switched, and half-media were changed every other day, on Day 12, HPC started to float and can be harvested and transferred to another 6-well plate for macrophage culture, on Day 16, the total harvest HPC were 3 million, 1.5 million were saved for sample analysis, and the other half 1.5 million HPC were continued to culture into macrophage, on Day 23, 1.4 million immature macrophage were harvested for sampling analysis and on Day 37, another 1.4 million immature macrophage were harvested for sampling analysis, finally on Day 44, the rest of 1 million macrophage were harvested for sampling analysis (iMacrophage ST). Genomic DNAs were isolated with 740954.20 NGS Prep Kit for Barcode Libraries in pScribe (CloneTracker XP™, LNGS-300) was used for genomic library amplification and all the primers and reagents for the first and second amplifications and sequencing of the genomically integrated barcodes.

### Barcode Counting

For each barcoded sample, fastq files were inspected to determine the BC14/BC30 sequence present at the defined location within the sequenced read, using perfect matching. Number of unique barcodes matching the expected barcodes from CloneTracker were counted, as well as numbers derived from earlier cellular stages. The Shannon Index/Maximum/Evenness were calculated for each sample.

## Code availability

Code used in this study is available from the authors upon written request.

### iMacrophage differentiation

Macrophages were induced with two independent previously published protocols^9,10^. For the first method listed as Protocol A, the HPC induction followed manufacturing protocol, briefly, on Day - 1, iPSCs were plated at 10-20 times lower plating density onto matri-gel coated 6 well plate with 2 ml mTeSR media, on Day 0, mTeSR media was removed and added 2ml Media A, on Day 2, 1ml media was removed and replaced with 1ml fresh Media A, on Day3, all media was removed and 2 ml of Media B was added, on Day 5, 1ml of Media B was removed and 1ml fresh Media B was added, 1ml Media change was performed on Day 5,7,10, on Day 12, all cells were transferred to a PDL-coated plate and cells were fed with Media C [500 mL IMDM (Gibco), 10% defined FBS (Gibco), 5 mL Penicillin/Streptomycin (Gibco)), 20 ng/mL of hIL3 (Peprotech), 20 ng/mL of hGMCSF (Peprotech) and 20 ng/mL of hM-CSF (Peprotech)]. It took average 3-4 weeks for the macrophages to be mature. For the second method, briefly, from Day 0 - 4, 10,000 cell (50 ul) in Step 1 media [Essential 8 Flex, 10uM ROCK Inhibitor, 50ng/ml BMP-4, 20ng/ml SCF, 50ng/ml VEGF], on Day 4 embryo bodies were formed, these embryo bodies were transferred to a 0.1% gelatin coated T175 flask in media Step 2 [X-vivo (10639011, LONZA) or StemPro 34 (10639011, Gibco), 25ng/ml IL-3, 100ng/ml M-CSF, 2mM GlutaMAX and 55um ß-mercaptoethanol], and media was changed every 7 days until Day 18, on Day 18, floating precursors were transferred to 6 well plate, each well with around 1.2 million cells in Step 3 media (RPMI1640+10% FBS, 2mM GlutaMAX and 100ng/ml M-CSF), on Day 25, mature macrophage were used for further analysis.

### iMicroglia ST and iMicroglia differentiation

iMicroglia ST (Protocol C) were induced following manufacture instruction with combination of induction kit and maturation kit (#100-0019 and #100-0020, Stem Cell Technologies) and was based on previous publication^21^. Briefly, on Day 0, iPSCs growing in 6 well plate were added with STEMdiff^TM^ Microglia Differentiation (#100-0019) with half media addition every other day till Day 12. On Day 12, the cell suspension was collected and centrifuged at 300g for 5 mins. Cells were then resuspended in STEMdiff^TM^ Microglia Differentiation Media then were seeded into Matrigel (Corning) coated plates until Day 24 with media addition every other day. On Day 24, cells were collected and centrifuged at 300g for 5 mins. Cells were then resuspended in STEMdiff^TM^ Microglia Maturation Media (#100-0020, Stem Cell Technologies). From Day 28-34, cells started to mature into iMicroglia ST and became less proliferative.

iMicroglia were differentiated as previously described^8^ (Protocol D). Briefly, iPSCs were cultured in mTeSR1 (Stem Cell Technologies) media on Matrigel (Corning) coated 6-well plates (Corning). When iPSCs reach around 80% confluency, they were dissociated using ReleSR (Stem Cell technologies), collected in 15ml canonical tubes followed by centrifugation for 5mins at 300xg and counted using trypan blue (Thermo Fisher Scientific). Typically, 200,000 cells/well were resuspended in mTeSR1 in low adherence 6-well plates (Corning). For day 1 to day 10, cells were cultured in DM medium [50% IMDM (Thermo Fisher Scientific), 50% F12 (Thermo Fisher Scientific), ITSG-X 2% v/v (Thermo Fisher Scientific), L-ascorbic acid 2-Phosphate (64 ug/ml, Sigma), monothioglycerol (400mM, Sigma), Poly(vinyl) alcohol (PVA) (10mg/ml, Sigma), Glutamax (1X, Thermo Fisher Scientific), chemically-defined lipid concentrate (1X, Thermo Fisher Scientific) and non-essential amino acids (Thermo Fisher Scientific)]. At day 0, embryoid bodies (EB) were gently collected, centrifuged at 100xg and resuspended in DM medium supplemented with 1uM ROCK inhibitor, FGF2 (50 ng/ml, Thermo Fisher Scientific), BMP4 (50ng/ml, Thermo Fisher Scientific), Activin-A (12.5ng/ml, Thermo Fisher Scientific) and LiCl (2 mM, Sigma), then incubated at in hypoxic incubator for 48hs (5% O^2^, 5% CO^2^, 37°C). On day 2, cells were gently collected and the media changed to DM medium supplemented with FGF2 (50ng/ml, Thermo Fisher Scientific) and VEGF (50 ng/ml, PeproTech) and returned to the hypoxic incubator for another 48hrs. On day 4 cells were gently collected and media changed to DM medium supplemented with FGF2 (50ng/ml, Thermo Fisher Scientific), VEGF (50 ng/ml, PeproTech), TPO (50 ng/ml, PeproTech), SCF (10ng/ml, Thermo Fisher Scientific), IL6 (50ng/ml, PeproTech) and IL3 (10ng/ml, PeproTech) and incubated in normoxic incubator (20% O^2^, 5% CO^2^, 37°C). At day 6 and 8, 2 ml of day 4 media was added in each well. On day 10, cells were collected, counted using trypan blue and frozen in Bambanker (Nippon Genetics) in aliquots of 500,000-1,000,000 cells. To start iMicroglia differentiation, cells were thawed, washed 1x with PBS and plated at 100,000-200,000 cells per well in 6-well plate coated with matrigel in iMicroglia media [(DMEM/F12 (Thermo Fisher Scientific), ITS-G (2% v/v, Thermo Fisher Scientific), B27 (2% v/v, Thermo Fisher Scientific), N2 (0.5% v/v, Thermo Fisher Scientific), monothioglycerol (200 mM, Sigma), Glutamax (1X, Thermo Fisher Scientific), non-essential amino acids (1X, Thermo Fisher Scientific)] supplemented with M-CSF (25 ng/ml, PeproTech), IL-34 (10 ng/ml, PeproTech) and TGFB-1 (50ng/ml, PeproTech). Cells were fed every 2 days and replated at day 22. On day 30, cells were collected and replated in iMGL media supplemented with M-CSF (25 ng/ml, PeproTech), IL-34 (10 ng/ml, PeproTech), TGFB-1 (50ng/ml, PeproTech), CD200 (100ng/ml, VWR) and CX3CL1 (100ng/ml, PeproTech). Cells were used at day 40 for functional and transcriptomic assays.

### Bulk RNA-seq analysis

Total RNA was extracted from cell pellets for duplicate samples using the Monarch Total RNA Miniprep Kit (New England BioLabs). RNA quality and concentration were checked on Tapestation (Agilent) with RNA ScreenTapes (Agilent). Bulk RNA-seq libraries were prepared using the QuantSeq 3’ mRNA-seq Library Prep Kit FWD (Lexogen). Final libraries were quantified with Tapestation (Agilent) with D1000 HS ScreenTapes (Agilent) and Qubit dsDNA HS reagents (Thermo Fisher Scientific). Sequencing was performed on NextSeq 550 (Illumina) for single-end 100bp reads. Reads were first trimmed to remove the sequencing adapters using bbduk (https://sourceforge.net/projects/bbmap/) and then aligned to the hg38 reference human genome using STAR^36^. Reads mapped to the genome were then assigned to genomic features (genes) using feature counts and Rsubread package^37^. Differential expression analysis was performed by using the negative binomial distribution from the R package DESeq2^38^. Principle components analysis (PCA) was performed to visualize the variance in the samples. PCA plots and volcano plots showing differentially expressed genes were plotted using the ggplot2 package (https://tidyverse.github.io/ggplot2-docs/authors.html). P-value and log fold change cutoffs used to filter the differentially expressed gene lists were 0.0001 and +/-1 respectively.

### Cytokine Profiling

300k/well macrophage of Wild Type and *SAMHD1* KO were cultured with Media C [500 mL IMDM (Gibco), 10% defined FBS (Gibco), 5 mL Penicillin/Streptomycin (Gibco)), 20 ng/mL of hIL3 (Peprotech), 20 ng/mL of hGMCSF (Peprotech) and 20 ng/mL of hM-CSF (Peprotech)]. 100ng/ml LPS was added to both WT and *SAMHD1* KO macrophage. 50ul of culture media of WT and WT treated with LPS and *SAMHD1* KO as well as *SAMHD1* KO treated with LPS were harvested, flash frozen and assayed for 48-plex protein levels by quantitative immunoassay (Eve Technologies).

### Lentiviral production

Lenti-X cells were seeded at 18,000,000 cells per plate in 15 cm dishes in 20 mL media (DMEM, 10% FBS) and incubated overnight at 37C, 5% CO2. The next morning, 8 mg sgRNA library plasmid, 4 mg psPAX2 (Addgene #12260), 4 mg pMD2.G (Addgene #12259) and 80 mL lipofectamine2000 (Invitrogen) were mixed into 1 mL serum-free OptiMEM (GIBCO), vortexed and incubated for 20 min at RT and added to the cells. At 72 h post-transfection, supernatant was harvested, passed through 0.45 um filters (Millipore, Stericup) and aliquots were stored at 80C.

